# PGViS: Personal Genome Variant interpretation Score for lung cancer genomes

**DOI:** 10.64898/2026.08.01.742250

**Authors:** Pallavi Surana, Pratik Dutta, Paolo Boffetta, Ramana Davuluri

## Abstract

Inherited lung cancer risk arises from both protein-coding and non-coding germline variants, but the functional non-coding component is largely uncharacterized. Genome-wide association studies and polygenic risk scores identify tag variants, not causal ones. Neither resolves which regulatory element is perturbed. DNA foundation models such as DNABERT decode non-coding variant effects directly from sequence, without a large GWAS cohort. What is missing is a patient-level framework linking these predictions to population-level variant prevalence. We present PGViS (Personal Genome Variant interpretation Score), a statistical framework that quantifies individual non-coding germline regulatory risk in non–small cell lung cancer (NSCLC). PGViS integrates three variant-level signals: DNABERT-predicted disruption at transcription factor binding and splice sites, the cancer v/s reference alternate allele frequency shift, and a regulatory interaction term derived from cancer-to-reference allele frequency ratios. Each signal is weighted by cohort prevalence which are aggregated into a single ancestry-matched, reference-normalized score per patient. We applied PGViS to germline whole-genome sequencing from 1,102 TCGA and CPTAC patients, using the 1000 Genomes Project (n = 2,504 individuals) as the normal population reference. PGViS separated adenocarcinoma (AD) and squamous cell carcinoma from controls in European ancestry and East Asian AD. Genes at contributing loci were enriched for *PI3K-Akt, Wnt*, DNA damage response, and epithelial–mesenchymal transition programs. Smoking-stratified analysis concentrated this signal on canonical NSCLC driver pathways. PGViS is modular: it accommodates cohorts with broader ancestral representation and can adapt to other solid tumors, offering a cost-effective route to personal-genome risk assessment from germline variants alone.

## 1. Introduction

Deleterious genetic variants in certain genes are key determinants of heritable cancer risk, including lung cancer. While protein-coding variants have been extensively studied, non-coding variants are also crucial, with most variants occurring in the gene regulatory regions [1–3]. While the functional effects of a single variant have been intensively studied, the joint impact of multiple variants has not been explored due to the complexity or lack of data. Due to the low frequencies of highly deleterious variants, aggregating these rare variants into combined categories is necessary [4, 5].

Population-level studies, such as the AllOfUs [6], UK BioBank [7], TCGA [8] and CPTAC [9] facilitate studying the impact of non-coding variants at the population level [5, 10]. Whole genome sequencing (WGS) datasets are relevant as they provide a comprehensive and precise approach compared to molecular testing, imaging, and lung histopathology by allowing early detection of genetic mutations, predicting disease risk, and offering deep mechanistic insights, all while being non-invasive and highly scalable. Further, Tumor mutation burden is an emerging biomarker which has been shown to be a prognostic indicator of survival in Lung cancer [11].

Large language models trained on DNA sequence can predict variant effects around core regulatory elements, including core promoters, splice sites, and transcription factor binding regions. Supervised approaches are limited by the training data available: variant sets skew toward common and well-annotated cell types, leaving rare cell types underrepresented [3, 12]. DNA foundation models address this through self-supervised pre-training on large unlabeled genomic corpora, followed by fine-tuning for specific downstream tasks. This reduces dependence on scarce labeled data and lets the model use both local and long-range sequence context, improving generalization across genomic contexts and particularly in the poorly characterized non-coding genome. [12–15].

DNABERT [13] identifies regulatory patterns in non-coding sequence directly from its learned embeddings, and has been applied to predicting non-coding variant effects at splice sites and TF binding sites [16, 17]. This enables identification of high-impact variants contributing to clinical risk, interpreted alongside exposures such as smoking. Existing non-coding variant scores, including FunSeq2 [4] and cis-X [2], operate at the individual variant level. They rank variants but do not yield a per-patient quantity comparable across individuals, and do not accommodate clinical stratification. PGViS fills this gap. It retains model-derived estimates of regulatory disruption, adds cohort- and population-level allele frequency behavior, weights each variant by its prevalence in the cohort, and aggregates across the variable-sized variant set each individual carries to produce one comparable score.

We present the PGViS, a statistical framework that integrates genomic language model predictions with population-level allele frequency data and clinical covariates to produce a per-patient measure of inherited regulatory burden. Using WGS from lung adenocarcinoma (AD) and squamous cell carcinoma (SCC) across two cohorts (TCGA and CPTAC), we characterize how this burden differs between histological subtypes, between smokers and non-smokers, and relative to a cancer-free reference population. We then map contributing loci to genes and test which biological programs they are enriched for and find the most disruptive TF programs. PGViS is a prioritization and stratification layer, not a validated diagnostic as it nominates high-impact regulatory variants and individuals with elevated inherited burden for functional and clinical follow-up. Its modular design permits extension to additional cohorts, ancestries, and tumor types.

## 2. Methods

### 2.1. Datasets for Variant Calling

Whole-genome aligned BAM files were downloaded from the GDC Data Portal using gdc-client [18]. We used solid tissue normal or blood-derived normal samples; where both were available, we selected the sample with higher coverage. Primary tumors were matched to their respective normals after download. Total data volume across cohorts was ~350 TB, with WGS coverage of 10–25x for CPTAC and 25–150x for TCGA. Germline variants were called with GATK HaplotypeCaller (v4.0.5.1) in gVCF mode (--ERC GVCF) against GRCh38, using chromosome-specific intervals for parallelization. Per-chromosome gVCFs were joint-genotyped, then filtered with GATK VariantFiltration (QUAL < 30, DP < 10, QD < 2, FS > 60, MQ < 40). Only PASS variants were retained, and multiallelic sites were normalized with bcftools [19].

The datasets are analyzed cohort and cancer wise, and the number of samples is summarized in **Table S1**. We also analyze smokers and non-smokers separately as we observe smoking-related patterns across the cohorts (for those with the number of samples >= 50). We fetch Allele count (AC), allele number (AN), and allele frequency (AF) annotations for the overall datasets and smoking status stratified datasets to analyze smoking-dependent and independent variants.

### 2.2. Validation dataset

As an independent normal germline reference, we used high-coverage (30×) WGS data from the 1000 Genomes Project (1kGP) [4, 5] on GRCh38 (GATK HaplotypeCaller SNV/INDEL calls; EMBL-EBI: ftp.1000genomes.ebi.ac.uk/vol1/ftp/data_collections/1000G_2504_high_coverage/. All 3,202 whole genomes (26 populations, five superpopulations: AFR, AMR, EAS, EUR, SAS) were used as the test cohort. Raw 1kGP VCFs were intersected with the smoking-dependent and smoking-independent regulatory regions, variants were used for the cancer cohorts, converted to BED (vcf2bed), and merged with per-sample genotype calls. Variants present in at least 5% of the cohort were retained and processed through the identical pipeline as explained above.

### 2.3. Population allele frequency sources

Variants were annotated against dbSNP v155 [20], which aggregates submissions from major population genomics resources including 1kGP, gnomAD, TOPMed, ALFA, and dbGaP, and catalogs over 1.2 billion unique rs identifiers. Population allele frequencies were taken from dbSNP. Where multiple frequency annotations existed, a representative general-population frequency was selected by the following priority: TOPMed (n = 158,000), gnomAD (n = 71,702), 1000 Genomes (n = 2,504 across 26 populations), then dbGaP. For the 1kGP comparison, 1kGP allele frequencies were excluded from the PGViS calculation to avoid using the control set in score construction.

### 2.4. Functional variant annotation and prioritization: a DNABERT prediction module

This module predicts functional effects of short nucleotide variants overlapping TF and histone binding sites and splice donors and acceptors [16]. Pre-trained DNABERT was fine-tuned separately on each region class. Splice site models used exon-intron junctions from GENCODE (v40), extended ±45 bp to 90 bp donor and acceptor sequences [13]; binding site models used ENCODE ChIP-seq data [21, 22] consisting of 4,228 experiments across 667 TFs and 33 histone marks in 301 bp windows (±150 bp). Splice site models achieved >95% accuracy; 462 of 690 binding site models, achieved >85% accuracy. All models are available: huggingface:duttaprat/DeepVRegulome [16].

Variant coordinates from the VCF files were intersected with these annotations, retaining variants recurrent within the study cohorts and present in dbSNP. Reference and alternate alleles were scored with the corresponding fine-tuned model. A variant was classified as functionally disruptive when the posterior probability of the reference sequence *p*(*s*) exceeded 0.5 and that of the mutant sequence *p*(*s*′) fell below 0.5; these thresholds are configurable. Magnitude of disruption was quantified by two complementary measures: the change in predicted probability and the log odds ratio between wild-type and mutant sequences as defined below.

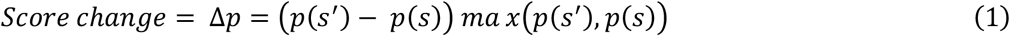

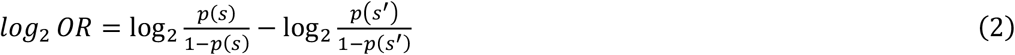

### 2.5. Variant Filtering and Functional Prioritization

Short germline variants (<10 bp) overlapping TFBS and splice regions were identified and filtered to enrich for functionally relevant short mutations among germline mutations in lung cancer.

1. Present in at least 5% of patients for smoking independent analysis. When stratifying by smoking status, an adaptive recurrence threshold was applied based on subgroup size: ≥50% (<100 samples), ≥40% (100–299), and ≥30% (≥300), with minimum counts of 40, 50, and 60.
2. Allele frequency shifts between cancer and matched normal samples from dbSNP v155 quantified using the change in alternate allele frequency (Δ*AF*_*alt*_ = *Can*_*altAF*_ − *Nor*_*altAF*_) and variants with |Δ*AF*_*alt*_| ≥ 0.1 were retained.
3. Allelic imbalance was further assessed using an absolute weighted imbalance metric (ΔWi), defined as the absolute difference in ref/alt ratios between cancer and normal, followed by inverse normal transformation (Δ*Wi*_*INT*_); variants with Δ*Wi*_*INT*_ ≥ 0 were selected.

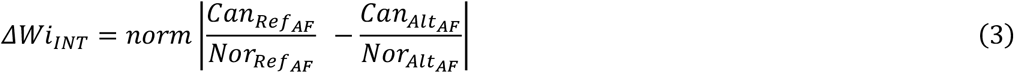
4. DNABERT fine-tuned model variant effect prediction module outputs threshold for variant independent analysis *p*(*s*)> 0.5 & *p*(*s*′)<= 0.5. For variant dependent analysis we use p(*s*)> 0.7 & *p*(*s*′)<= 0.3. This threshold summarizes only the loss of function mutations where the probability of the alternate allele is lesser than that of the reference allele.
5. We excluded variants with a minor allele frequency [23] < 0.05, because sample sizes are small and uneven across groups, leaving insufficient power to reliably detect associations.

### 2.6 Personal Genome Variant interpretation score calculation

Each patient carries a variable number of germline variants, each characterized by three features: the change in regulatory interaction strength (Δ*Wi*_*INT*_), the alternate allele frequency shift (Δ*AF*_*alt*_), and the DNABERT-derived *log*_2_ *OR*. Because variant counts differ across patients, naive aggregation would not yield comparable values. Each variant was therefore weighted by the ratio of samples carrying it to the maximum carrier count of any variant in the cohort. A feature-specific patient score was the mean weighted value across that patient’s variants.

Scores were computed separately for smoking-dependent and smoking-independent variant sets, retaining only variants shared between TCGA and CPTAC within a given context, and the two were averaged. Integrated features were standardized by a robust, ancestry-matched z-score, using the median and median absolute deviation among 1kGP controls of the same superpopulation rather than cohort-wide mean and standard deviation, then rescaled to unit variance so no feature dominated by numeric range. Feature weights were derived by principal component analysis, with absolute PC1 loadings normalized to sum to unity. PGViS for patient *p* is the weighted sum of the three standardized features (K = 3), as given below.

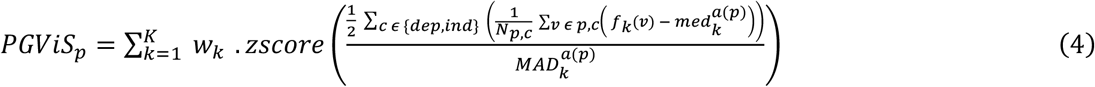

Here, *f*_k_(*v*) is the value of feature *k* for variant *v*, and *c* indexes the smoking-dependent and smoking-independent variant sets, with *N*_p_,*c* the number of variants carried by patient *p* in context *c*. The term *a*(*p*) denotes the superpopulation assigned to patient *p* (AFR, EAS, or EUR), and 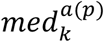 and 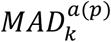 are the median and median absolute deviation of feature *k* across 1000 Genomes control individuals of that superpopulation. The function zscore(·) denotes standardization to zero mean and unit variance across all individuals in the study; the resulting score is unbounded and centered at zero and is not scaled to a fixed interval. Finally, *w*_k_ is the weight of feature *k*, given by its absolute loading on the first principal component, normalized so that weights sum to one.

### 2.7. Functional interpretation of variant-associated regions

Germline variants affecting TF binding functional sites were merged into independent genomic loci (868 smoking-dependent, 367 smoking-independent) and mapped to genes using rGREAT v2.10.0 under the hg38 basal-plus-extension model (−5/+1 kb basal domain, bidirectionally extended up to 1 Mb; assembly gaps excluded) [24]. Mapped genes were resolved to HGNC symbols, re-linked to original window coordinates by interval overlap, and compiled for downstream pathway analysis.

Pathway analysis was conducted using cGSA [25], an LLM framework that partitions a STRING interaction network into EdMot communities to prevent hub-gene inflation, screens candidate Enrichr terms (p < 0.001), and selects representative pathways per community. Using GPT-4o (v20240513) as the backend model, terms were assigned relevance scores (0.0–10.0; thresholds: 5.5 moderate, 7.0 high) based on the specified experimental context (smoking-related and overall analysis) and research objective (non-small cell lung cancer). We also analyze the top TFs disrupted, their prevalence in smoking related and unrelated cohorts and their mutations enriched pointing to identify key regulatory TFs and their regions disrupted in NSCLC.

## 3. Results

### 3.1 Framework to identify variants with a functional impact in NSCLC

The analysis focuses on finding functionally relevant short variants in the non-coding regulatory regions, including splice acceptor/donor sites and transcription factor (TF) binding regions near transcription start sites, where sequence alterations can influence gene regulation. We focus on germline mutations to then figure out the risk conferred by them to the pathogenicity of NSCLC.

WGS variants from CPTAC and TCGA cohorts (AD, SCC) are first evaluated using DNABERT-based fine-tuned models to quantify sequence-level regulatory impact. In parallel, variant frequency information across patients is summarized to capture population-level enrichment and distributional properties. These components are combined to generate variant–patient pair scores that reflect both predicted functional impact from transformer-based attention architecture and observed changes in allele frequency in the normal populations (dbSNP) and in cancer cohort (**Fig. 1**).

**Fig. 1:**
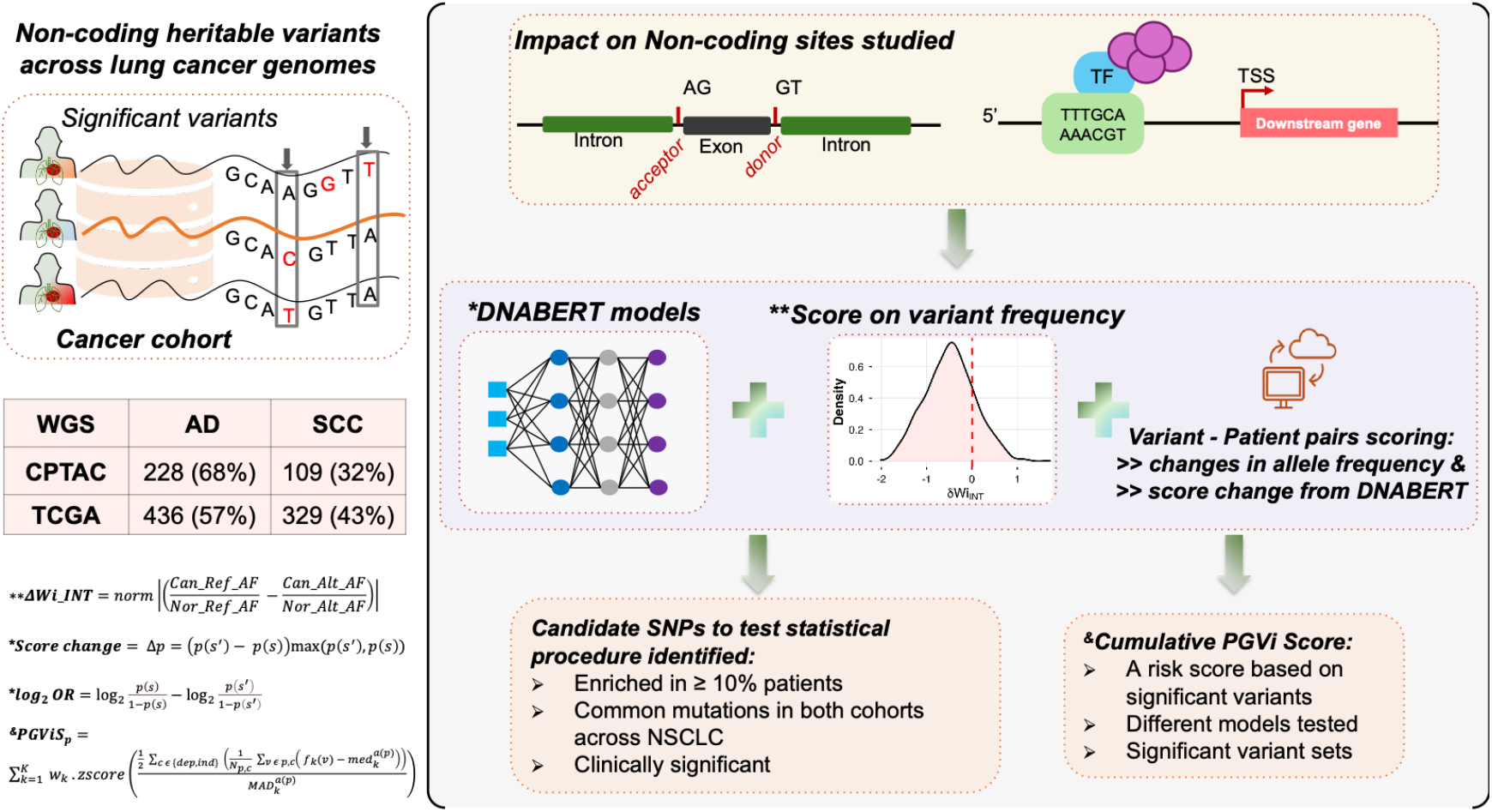
Framework for PGViS. Non-coding germline variants from WGS datasets for 2 cohorts are analyzed (TCGA, CPTAC). The DNABERT models are used to provide the variant impact information based on the TF binding sites and splice sites it affects. Allele frequency based Δ*Wi*_*INT*_ is calculated based on the ratio of cancer and normal allele frequencies. All this information is used to then develop a cumulative risk score for each sample/patient of NSCLC.

We identify a pool of germline mutations after basic set of filtering as written in **Methods 2.1**. In the AD cohort, TCGA contains ~16.43M unique functional sites mapping to ~8.03M dbSNP-annotated variants, while CPTAC includes ~15.56M unique functional sites mapping to ~6.03M dbSNP variants. One variant can affect multiple functional sites in proximity owing to the higher number of them in this table. Their overlap yields ~13.08M shared functional sites, mapping to

~3.98M unique variants (**Table S2**). The scope of this study only focuses on variants common between dbSNP and the cancer cohort. In SCC cohort we have ~3.3M common variants across the 2 cohorts and ~11.97M unique functional sites. Overall, AD shows higher variant and functional site counts than SCC, and the TCGA–CPTAC overlap defines a large, high-confidence, dbSNP-validated non-coding variant set used for downstream analyses (**Table S3A-B)**.

The thresholds to get the final list of candidate SNPs are 5 steps which include short variants (<10 bp) were selected for analysis if they were present in at least 5% of patients and showed a substantial change in allele frequency (|Δ*AF*_*alt*_| ≥ 0.1). Variants were further required to have a non-negative DNABERT-derived Δ*Wi*_*INT*_ (Δ*Wi*_*INT*_ ≥ 0), high reference sequence confidence (ref_prob > 0.7), and low alternate sequence confidence (alt_prob ≤ 0.3), indicating a predicted functional impact. Rare variants in cancer cohort were excluded by filtering out mutations with minor allele frequency [23] < 0.05. This is done as we cannot account for them due to the smaller sample sizes hence not enough power in the analysis to predict impact of rare variants. Only variants common across cohorts were retained for downstream analyses. The last threshold used is the *log*_2_ *OR* from DNABERT-finetuned models where we use a valley point cutoff based on the distribution of the significant mutations to determine the success of variant disruption in the right end of the plot (**Fig. S1**). The patterns differ between smoking dependent and independent analysis.

Candidate SNPs are then prioritized using statistical criteria, including enrichment in at least 5% of patients, presence across both cohorts, and clinical relevance. Finally, significant variants are aggregated at the patient level to compute a cumulative risk score. Multiple model configurations and variant subsets are evaluated, and only statistically significant variant sets are retained, yielding a robust PGViS score that integrates regulatory impact and cohort-level mutation patterns. **Table S3** shows the summary of thresholds used to arrive at final list of significant mutations used further in the analysis. This analysis is done combining all samples in the cohort without classification further based on clinical features. We get ~1-1.5k significant mutations based on the threshold across the TCGA and CPTAC cohorts that we use in further analysis.

We next examined variant overlap between the TCGA and CPTAC cohorts. More variants from TCGA contribute to the shared set than from CPTAC. However, CPTAC yields a greater number of significant variants in both AD and SCC. This likely reflects CPTAC’s smaller sample size and lower sequencing depth (**Table S4-top**). The shared and cohort-specific variants may point to common and distinct risk patterns across patient groups.

### 3.2 Smoking-Dependent Variant Identification in TCGA and CPTAC

Variant distributions stratified by smoking history differed across cohorts and ancestries. In the CPTAC East Asian (EAS) AD cohort, smoker and non-smoker distributions largely overlap, indicating that smoking status does not strongly differentiate variant burden in this subgroup possibly reflecting biological differences, exposure patterns, or limited sample size (**Fig. 2A**). In contrast, European (EUR) patients show clear separation in both AD and SCC, with smokers carrying higher and more dispersed variant counts. AFR and unknown-ancestry groups show weaker or inconsistent separation, likely due to smaller cohort sizes (**Fig. 2B**). Separation is sharpest in TCGA, particularly for SCC (**Fig. S3**).

**Fig. 2:**
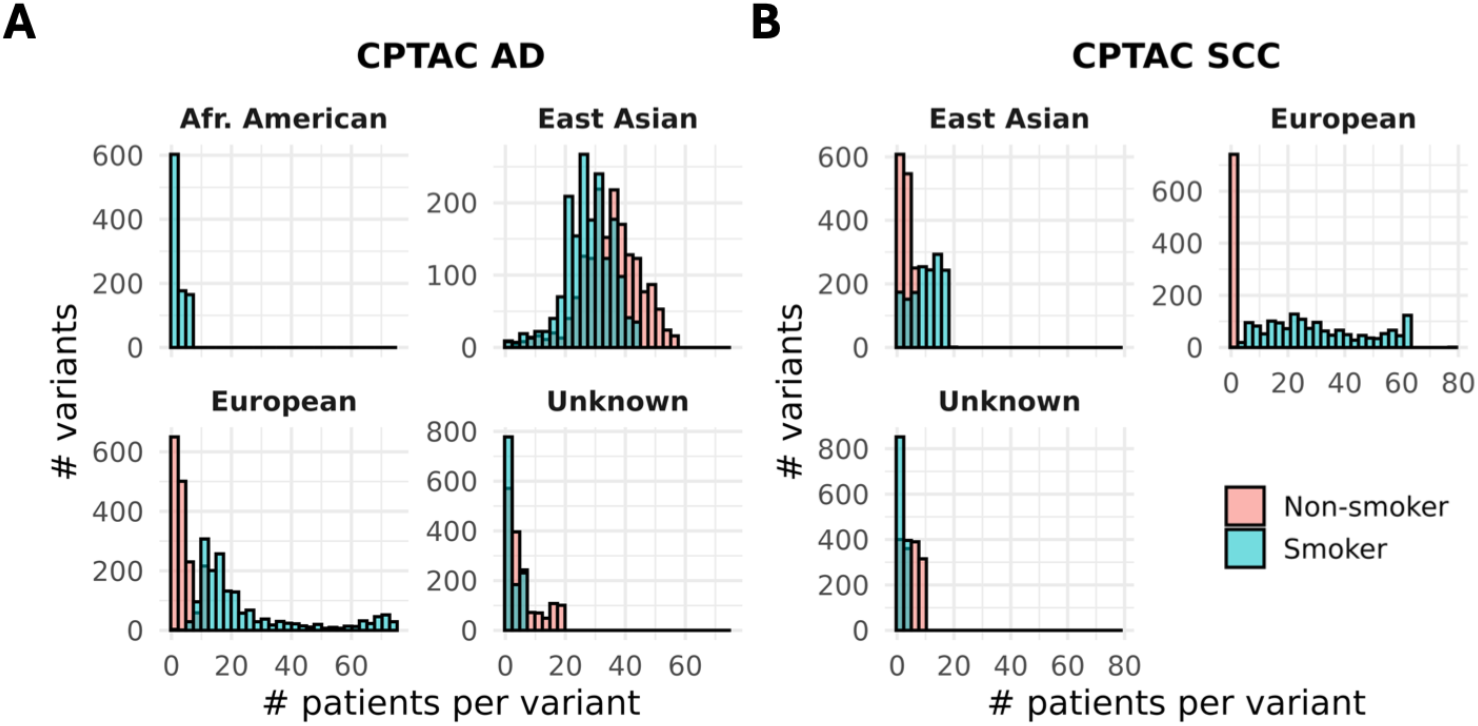
Variant frequency spectra in the CPTAC cohort, stratified by ancestry and smoking status, for (A) AD and (B) SCC. Each histogram shows the number of variants (y-axis) carried by a given number of patients (x-axis). EAS AD shows near-complete overlap between smokers and non-smokers, whereas EUR shows divergent spectra in both AD and SCC. AFR and Unknown-ancestry groups are shown for completeness but are too small to interpret.

This heterogeneity motivates stratifying by smoking status. Separate analysis captures smoking-dependent signatures where present, as in EUR, while avoiding signal dilution where smoking status has little effect on variant burden, as in EAS AD. We do not stratify by ancestry because of insufficient sample size. CPTAC reports ~3.5× more significant variants among smokers than TCGA, and ~10,000 in non-smoker AD (**Table S4-bottom**).

### 3.3 Allele frequency dependent score bias in cohorts

Having defined smoking-independent (all samples analyzed together) and smoking-dependent (smokers and non-smokers analyzed separately) sets of significant variants, we next asked whether the Δ*Wi*_*INT*_ score separates AD from SCC (**Table S5**). Before interpreting these distributions, we note a systematic feature of the variant sets themselves: the number of significant variants scales inversely with cohort size. The smallest cohorts yield the largest variant sets (CPTAC AD non-smoker, n = 87, 18,266 unique functional sites), while the largest yield the fewest (TCGA All AD, n = 434, 1,363 sites). Because significance here depends on allele frequency contrasts, smaller cohorts produce noisier frequency estimates and therefore more variants passing threshold, so variant set size reflects cohort size at least as much as underlying biology.

With that caveat, **Fig. 3** compares the distribution of Δ*Wi*_*INT*_ across cancer histologies and smoking status in the CPTAC and TCGA cohorts. In the CPTAC cohort, Δ*Wi*_*INT*_ shows a clear separation between AD and SCC: AD variants concentrate sharply near zero, while SCC variants show a broader distribution shifted toward higher values, indicating that Δ*Wi*_*INT*_ captures histology-associated differences in variant impact in this cohort (**Fig. 3A**). In TCGA, the AD and SCC distributions overlap almost completely across the same range, giving no useful discrimination between histologies. When stratified by smoking status, CPTAC SCC smokers show a shift in Δ*Wi*_*INT*_ relative to AD smokers, with a secondary mode near 0.6 that is more distinct in SCC, whereas the corresponding TCGA distributions remain largely superimposed (**Fig. 3B**).

**Fig. 3:**
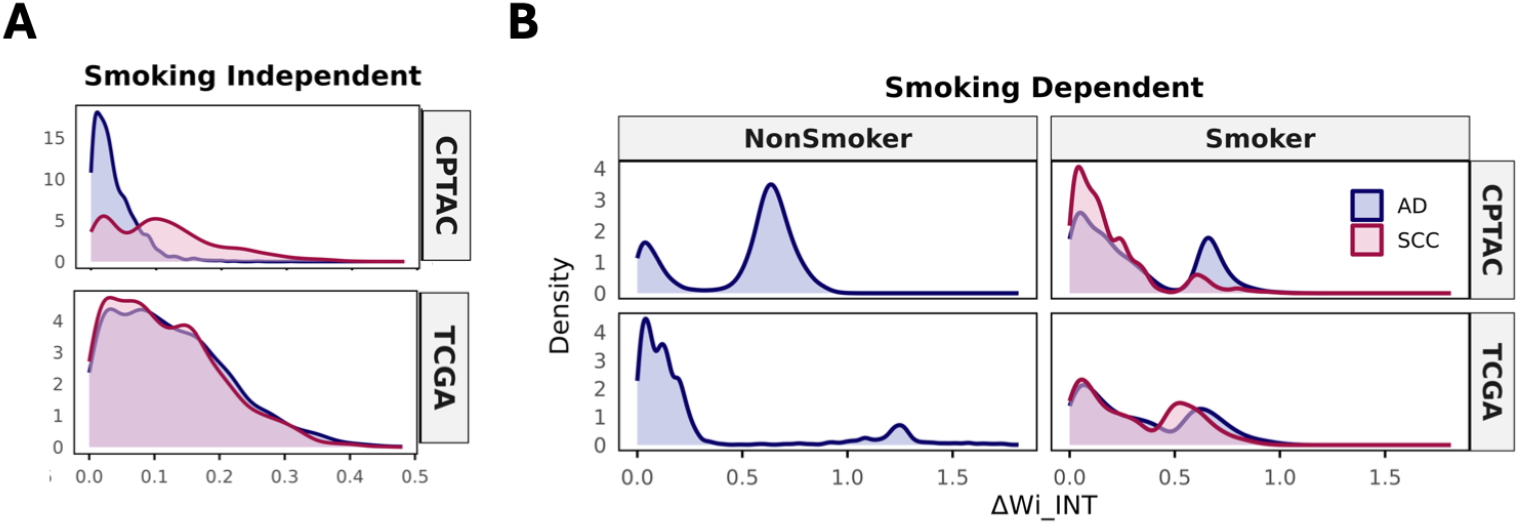
Distribution of Δ*Wi*_*INT*_ (A) distinction between AD and SCC in CPTAC cohort but not TCGA, (B) Smokers in SCC show clear distinction from AD in smokers across CPTAC but not as profound in TCGA cohort.

These results indicate that Δ*Wi*_*INT*_ reflects histology- and smoking-associated variant effects more strongly in CPTAC than in TCGA. However, the CPTAC variant sets are 3–8 times larger than the TCGA sets and are enriched for lower-frequency variants, so the cohort difference in separation cannot be cleanly attributed to biology rather than to variant set composition. We therefore treat Δ*Wi*_*INT*_ as one contributing component of the PGViS score rather than as a standalone discriminator, and retain cohort-specific, smoking-stratified analysis in subsequent modeling.

### 3.4 PGVi Score separates Non-small cell lung cancer types from Control cohort

To derive a patient-level measure of non-coding germline risk, we integrated variant-level signals of regulatory disruption and population enrichment into a single score. We first assessed whether candidate components contributed complementary information (**Fig. S3A**). Most pairwise correlations were near zero in both smoking-stratified and pooled sets, indicating that each captures a distinct dimension of variant impact. The exception was a reproducible moderate negative correlation between log_2_ OR and ΔAF_alt_ (r = −0.41 smoke-split, −0.49 pooled); we retained log_2_OR and dropped ΔAF_alt_. The final score comprises *log*_2_ *OR*, Δ*Wi*_*INT*_ and Δ*AF*_*alt*_ (the difference between cancer and dbSNP alternate allele frequencies), each weighted by cohort-level representation and aggregated across all qualifying variants carried by a patient. Stability of this correlation structure across smoking strata indicates that the internal geometry of the score is not driven by smoking status.

We next asked whether PGViS separates cancer patients from a cancer-free reference. We scored 1kGP through the identical pipeline, with standardization parameters fixed on the cancer cohorts. Because two of three components are allele-frequency-derived and allele frequencies vary with ancestry independent of disease status, we restricted this comparison to EUR-ancestry samples (**Fig. 4A–B**). Distributions were near-completely separated in all four cohort-subtype comparisons: AD versus 1kGP gave Cohen’s d = 6.36 (CPTAC) and 5.33 (TCGA); SCC versus 1kGP gave 6.14 and 3.72. All permutation p < 0.001. Cancer samples shifted toward higher PGViS in every comparison, consistently across two independently generated cohorts. The pattern held in EAS-ancestry samples, where PGViS separated CPTAC AD from 1kGP with d = 4.41 (p < 0.001; **Fig. S3B**).

**Fig. 4:**
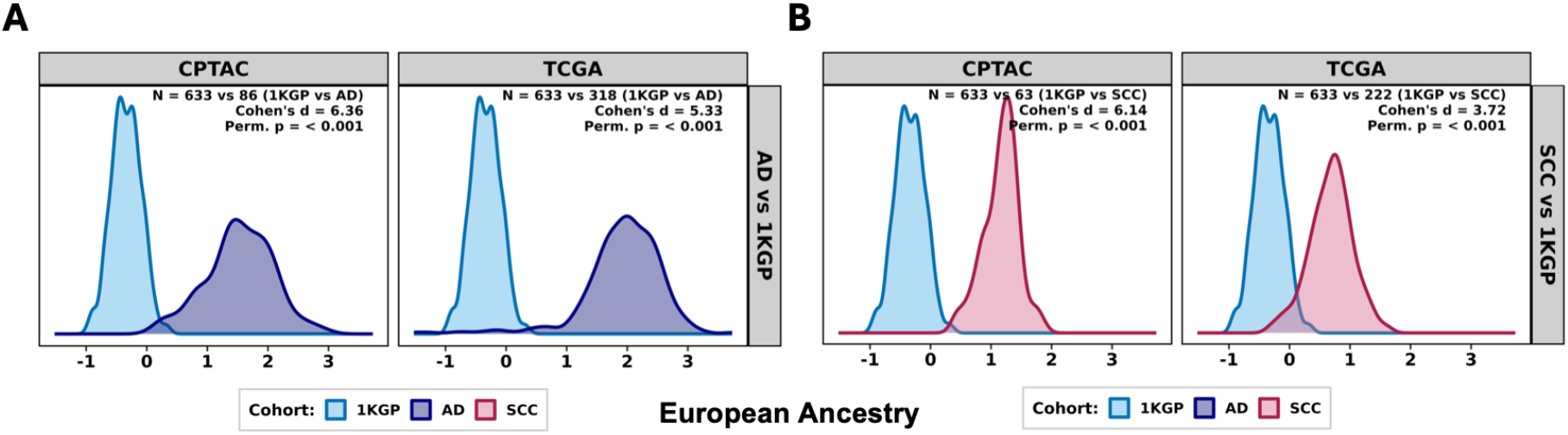
PGViS score construction and validation against a normal reference. (A) Pairwise Pearson correlations among candidate variant-level metrics, shown for the smoking-stratified sets (top) and all samples (bottom). (B) PGViS score distributions for AD and SCC patients versus the 1kGP cancer-free reference, restricted to European (EUR) ancestry and shown separately for CPTAC and TCGA. Sample sizes, Cohen’s d, and permutation p-values are annotated per panel.

PGViS is computed by assigning each germline variant a composite impact score integrating regulatory interaction change, predicted functional effect, and allele frequency shift, weighted by cohort-level representation, then aggregating across all qualifying variants carried by a patient to give a single continuous measure of cumulative non-coding burden. By combining largely independent regulatory signals, PGViS avoids reliance on any single metric.

### 3.5 Variant-mapped genes enrich for growth-factor signaling, genome maintenance and EMT pathway

In the smoking-stratified set, PI3K–Akt signaling scored highest (9.0) and formed the largest community (70 genes), containing canonical NSCLC drivers and receptor tyrosine kinases (*PIK3CA, ERBB3, FGFR3, PDGFRA, KIT, HGF, CCND1*). Four communities reached 8.5: Wnt signaling (*AXIN2, WNT5A, SFRP2, SOX9, TCF4, MYCN*), RNA metabolism and the ubiquitin– proteasome system (*PSMC4, PSMD7, USP12/13/14, HNRNPD*), DNA damage response and repair (*PRKDC, RAD52, RAD17, RNF168*), and epithelial–mesenchymal transition (*COL5A2, LAMA5, MXRA5, FSTL1*). RUNX transcription, Rho GTPase signaling (*ARHGEF12, DLC1, RHOB, PREX1*), and oxidative phosphorylation (*CYC1, UQCRFS1, ISCU, CS*) also scored highly (**Fig. 5A**).

**Fig. 5:**
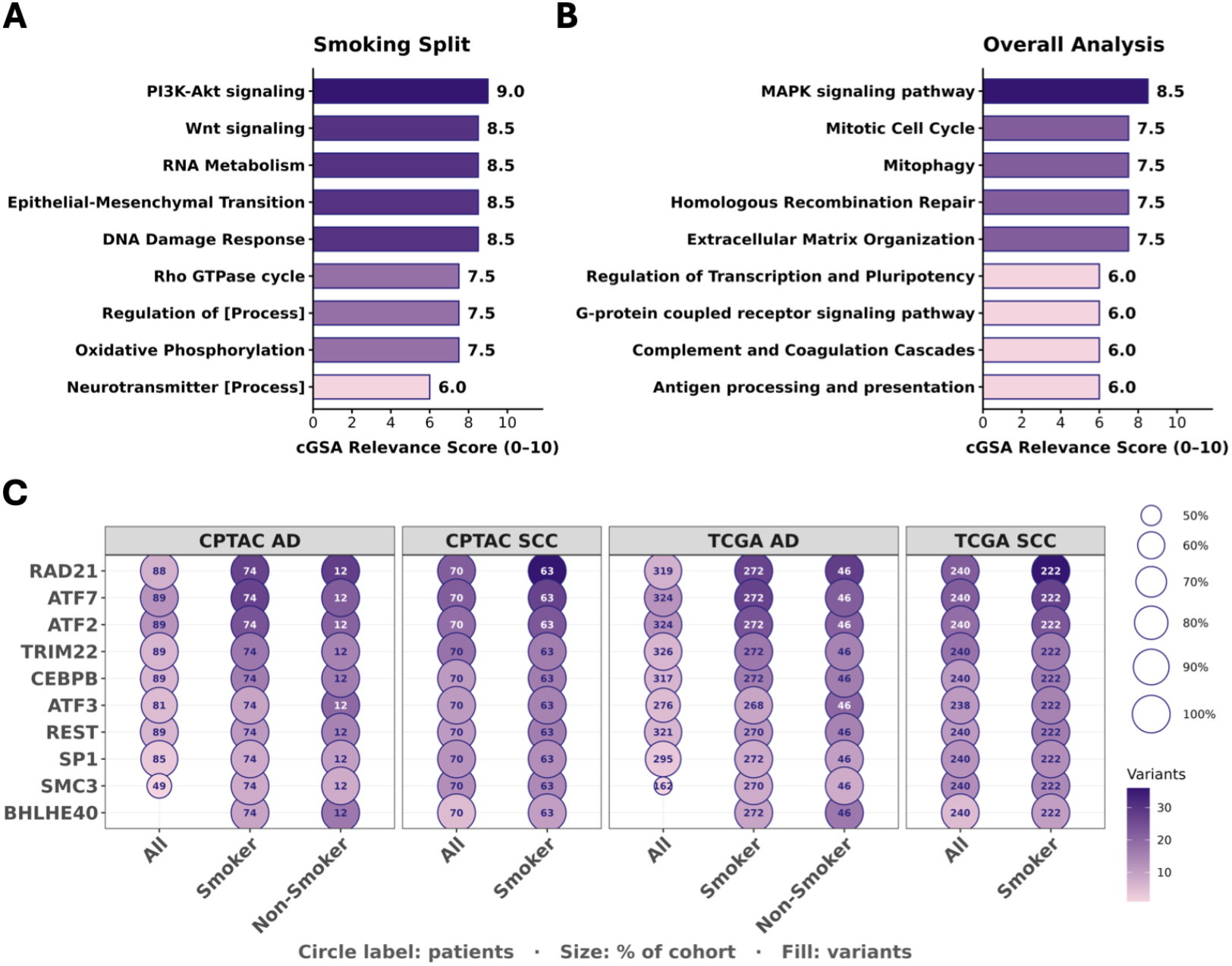
Contextual pathway relevance and recurrently disrupted transcription factors. (A–B) Candidate pathways scored by the cGSA framework (GPT-4o backend) for the smoking-stratified (A) and unstratified (B) analyses. Bars show contextual relevance on a 0–10 scale; scores > 7.0 denote high relevance. (C) Ten most frequently disrupted TFs across cohorts, histologies, and smoking strata. Rows are TFs ranked by patient recurrence; columns are strata (All, Smoker, Non-Smoker) within each cohort–histology panel. Circle labels give patient counts, circle size the percentage within cohort, and fill intensity the number of distinct disrupted variants. Non-smoker SCC strata are omitted for insufficient sample size. Per-stratum totals are in **Table S6**.

Pooling smokers and non-smokers recovered a narrower, less driver-centric signal. MAPK signaling was the sole community above 8.0 (8.5; *MAP2K3, MAP2K6, MAPK13, MAPK3, RPS6KA1, HSPB1*), followed at 7.5 by mitotic cell cycle, mitophagy, extracellular matrix organization (*MMP2, TIMP3, COL5A1, ADAMTS2*), and homologous recombination repair (*PRKDC, ERCC4, XRCC3*). Remaining communities scored 6.0, with cGSA attributing these to indirect links through immune evasion or transcriptional plasticity. Modeling smoking status therefore shifts the signal toward canonical NSCLC driver programs (**Fig. 5B**).

At the level of individual regulators, top ten transcription factors were disrupted in nearly every patient across both cohorts, both histologies, and all smoking strata: *RAD21, ATF7, ATF2, TRIM22, CEBPB, ATF3, REST, SP1, SMC3*, and *BHLHE40* (**Fig. 5C**), each affected in ≥90% of patients in nearly all strata. Recurrence was not histology-driven; AD and SCC converged on similar regulators. Two exceptions track the pathway-level dilution seen on pooling: *SMC3* fell to roughly half of patients in unstratified AD (49/89 CPTAC; 162/319 TCGA), and *BHLHE40* was absent.

These factors fall into three groups with independent support in lung cancer. The AP-1 family (*ATF2, ATF3, ATF7*) is elevated in lung tumors, promotes invasion, and is linked to shorter survival, with *ATF3* induced by cigarette smoke in airway epithelium [26, 27]. Cohesin subunits (*RAD21, SMC3*) are amplified in AD, where *RAD21* predicts poor prognosis [28]. The remainder are stress-responsive or broadly acting: *CEBPB, SP1, TRIM22, BHLHE40*, and *REST* are each altered in lung tumors [29–31]. This convergence on architectural and stress-response regulators rather than lineage-restricted factors is consistent with reporter screens showing that most lung cancer risk loci act through allele-specific binding at multi-locus factors [32, 33]. Such shared regulatory substrate provides the background on which the histology-specific and smoking-specific pathway programmes described above are superimposed.

## 5. Discussion

Within the non-coding genome, evolutionarily constrained regions are enriched for functional regulatory elements and disease-associated variation, providing a principled link between biological annotation, disease association, and natural selection. This motivates risk modeling approaches that prioritize non-coding variants not only by their genomic location but also by their predicted regulatory impact.

PGViS provides an extensible framework for translating heterogeneous sets of non-coding germline variants into an interpretable, patient-level risk score for non–small cell lung cancer. A key strength of this approach is that it explicitly combines complementary information sources: (i) sequence-based functional disruption predicted by DNABERT models in regulatory contexts (log_2_OR TFBS and splice regions) and (ii) cohort- and population-level allele frequency behavior captured by ΔAF_alt_ and ΔWi_INT_. By design, this integrates “what a variant is predicted to do” with “how that variant behaves across individuals,” allowing PGViS to summarize risk-relevant regulatory perturbations that would be missed when variants are considered in isolation or when analysis is restricted to coding regions. The weak correlations among the retained score components indicate that PGViS benefits from integrating non-redundant signals rather than amplifying a single dominant metric.

The separation of PGViS distributions between cancer patients and the 1kGP reference, together with the differences observed between adenocarcinoma and squamous cell carcinoma, supports the premise that inherited regulatory burden differs both from the general population and across histological subtypes. This is consistent with the broader understanding that AD and SCC arise through partially distinct etiologic and molecular trajectories, and that germline variation may influence these trajectories through regulatory control rather than protein alteration alone. PGViS captures these differences using only germline non-coding variants filtered to functional regions, and the pattern is reproduced in two independently generated cohorts and in two ancestry groups, which argues against a signal specific to any single dataset. The smoking-stratified analysis further indicates that smoking history modulates the detectable variant landscape in a subgroup-dependent manner, reinforcing the need to model clinical context when interpreting inherited variation in lung cancer.

A practical contribution of PGViS is its aggregation strategy for unequal-sized bags of variants per patient. Weighting each variant by its cohort prevalence reduces sensitivity to extremely rare events that cannot be estimated robustly at moderate sample sizes, while allowing recurrent disruptive variants to contribute proportionally. Patient-level averaging and ancestry-matched standardization then place patients on a comparable scale across features with different units and dynamic ranges, enabling downstream analyses such as subtype discrimination and risk stratification.

Mapping the variant-associated loci to genes situates the prioritized variants in interpretable biology. The recovered communities were dominated by growth-factor and developmental signalling (PI3K–Akt, Wnt, MAPK), genome maintenance (DNA damage response, homologous recombination repair), and programmes associated with invasion and tumour–stroma interaction (epithelial–mesenchymal transition, extracellular matrix organisation, Rho GTPase signalling): processes central to NSCLC biology and, notably, recovered here from germline regulatory variants rather than somatic drivers. That the smoking-stratified gene set scored consistently higher and yielded more driver-proximal pathways than the unstratified set parallels the variant-level results and supports stratification as more than a technical convenience. At the regulator level, the same small set of transcription factors was disrupted in nearly every patient regardless of cohort, histology or smoking status, and these were dominated by architectural and stress-response factors (cohesin, *AP-1, CEBPB, BHLHE40*) rather than lineage-restricted ones. This suggests that inherited regulatory burden in NSCLC concentrates on broadly acting regulators, which provide a shared biology of TF disruption on which the histology- and smoking-specific pathway programmes described above are superimposed. These pathway results should be read as descriptive context for the variant sets rather than as independent evidence, for reasons given below.

Several limitations should be considered. DNABERT predictions are a scalable but model-derived proxy for regulatory impact, and may miss long-range chromatin interactions, cell-type specificity absent from training data, or context-dependent effect directionality. Our filtering emphasizes recurrent variants and excludes low-frequency ones for lack of power, which may under-represent highly penetrant rare alleles; variant set size also scales inversely with cohort size, so composition partly reflects sample size rather than biology. PGViS has further not been evaluated predictively ie., all comparisons contrast prevalent cases against a reference at a single time point, and no discrimination or calibration metrics are reported.

PGViS is intended as a foundation for broader personalization of germline risk modeling. Immediate next steps include evaluating predictive performance in external cohorts processed under a uniform pipeline, testing alternative aggregation functions such as robust means or learned nonlinear combinations, and integrating orthogonal signals such as sequence conservation, epigenomic accessibility, eQTL and allele-specific expression evidence, or polygenic risk scores. Clinically, PGViS could serve as a prioritization layer to flag individuals with elevated inherited regulatory burden and to nominate high-impact variants for functional follow-up. More broadly, the framework illustrates how genomic language models can be coupled with population genetics and clinical stratification to move from variant-centric interpretation toward patient-level risk quantification in the non-coding genome.

## Acknowledgements

This work was supported by:

- National Institutes of Health, National Library of Medicine [R01LM013722 to R.D.]
- Partial by NIH T32 Predoctoral training grant [T32GM148331 to P.S.]
- Davuluri lab and computing cluster at Stony Brook University, Medicine.

Preprint of an article submitted for consideration in Pacific Symposium on Biocomputing © [2027] World Scientific Publishing Co., Singapore, http://psb.stanford.edu/.

## Code Availability

The code will be made available once the manuscript is accepted.

