## Supplementary Information for "PGViS: Personal Genome Variant interpretation Score for lung cancer genomes"

### Supplementary Information for Surana et. al, PGViS

#### Supplementary Figures

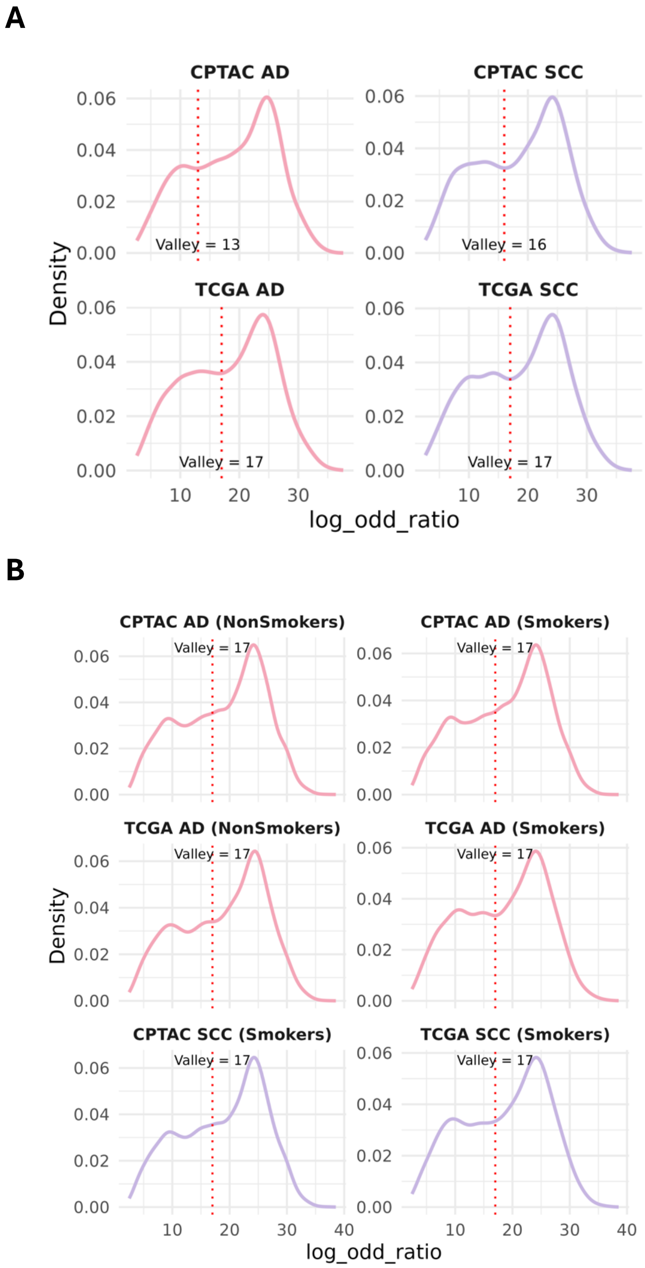


**Fig. S1:** Density plot of the DNABERT-derived $log_2\, OR$ for all analyzed variants. The distribution shows an approximately bimodal pattern, indicating two distinct groups of variant effects. An initial DNABERT LOR cutoff was applied to identify potentially significant mutations, followed by additional thresholding to refine the final mutation set. The right-hand peak of the distribution, corresponding to variants with higher predicted regulatory impact, was used as a key threshold for selecting significant mutations included in downstream analyses.

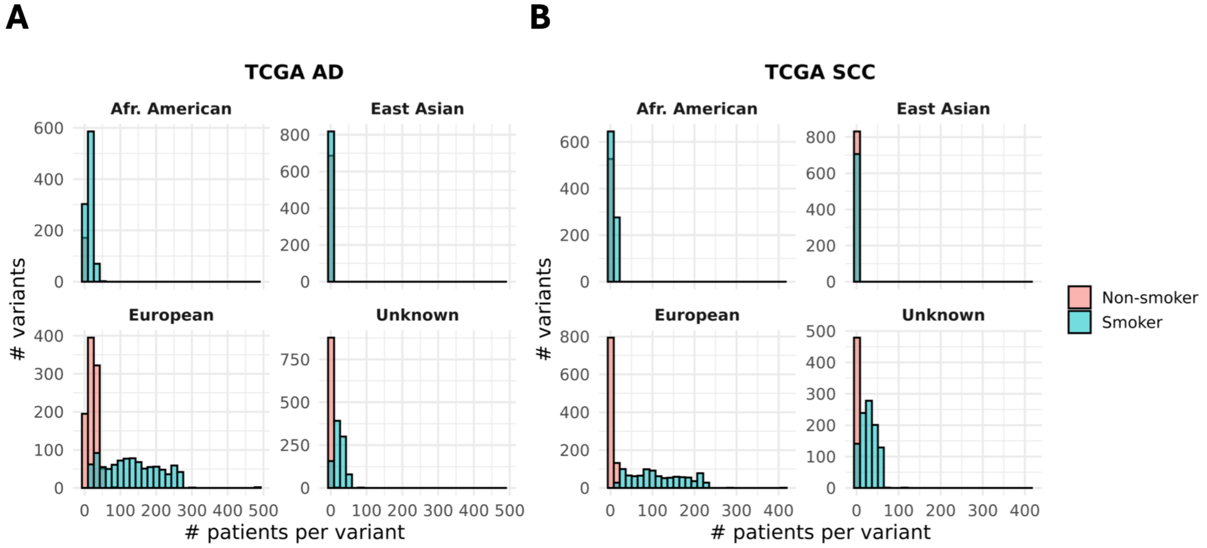


**Fig. S2:** Distribution of variant burden per patient across ancestry groups in TCGA cohort (A) AD and (B) SCC stratified by smoking status shows clear distinction between smokers and nonsmokers across ancestries.

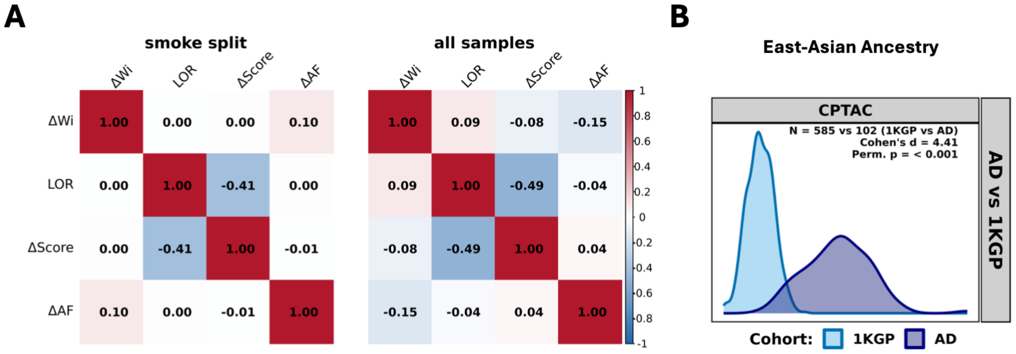


**Fig. S3:** PGViS score distributions for EAS-ancestry samples. CPTAC AD patients (n = 102) versus the 1kGP EAS reference (n = 585). Cohen's d, and permutation p-value are annotated. TCGA and SCC comparisons are not shown for EAS owing to insufficient sample size. *Note*: ΔScore is alternatively used as $\Delta AF_{alt}$

#### Supplementary Tables

**Table S1.** Number of samples in AD and SCC of lung across CPTAC and TCGA cohorts. The % is with respect to the total samples in each cohort.

| WGS dataset | Lung AD | | | | Lung SCC | | | | Total Across |
|---|---|---|---|---|---|---|---|---|---|
| | Smoker | NoSmoker | Unknown | Total | Smoker | NoSmoker | Unknown | Total | |
| **CPTAC** | 133 | 87 | 8 | 228 (68%) | 84 | 16 | 9 | 109 (32%) | 337 |
| **TCGA** | 362 | 61 | 13 | 436 (57%) | 304 | 14 | 11 | 329 (43%) | 765 |
| **Total** | 495 | 148 | 21 | 664 | 388 | 30 | 20 | 438 | 1102 |

**Table S2:** Table summarizing those variants where the ref and alt alleles as the positions match between cohort (CPTAC, TCGA) and dbSNP v155. This is the starting group of variants across functional sites used in this study.

| Cancer: | Cohort | Number of sites across TFBS + Splice Region | | |
|---|---|---|---|---|
| | | Unique Functional Site | # Variants | dbSNP IDs |
| **ADD** | TCGA: all variants | 16431811 | 8080612 | 8034254 |
| | CPTAC | 15555095 | 6057649 | 6031724 |
| | TCGA ∩ CPTAC | 13081924 | 3980558 | 3969128 |
| **SCC** | TCGA: all variants | 15971353 | 6949822 | 6914833 |
| | CPTAC | 13850252 | 4383867 | 4369786 |
| | TCGA ∩ CPTAC | 11974579 | 3334336 | 3326272 |

**Table S3:** Summary of thresholds used to arrive at final list of significant mutations used further in the analysis. This analysis is done combining all samples in the cohort without classification further based on clinical features. (A) Thresholds used to get significant variants across functional sites for both AD and SCC in the CPTAC cohort. (B) Thresholds used to get significant variants across functional sites for both AD and SCC NSCLCs in the TCGA cohort.

**A**

| **Cancer** | **Thresholds** | **Significant # of across TFBS + Splice Region** | | |
|---|---|---|---|---|
| | | **Unique Functional Site** | **CPTAC Variants** | **dbSNP IDs** |
| **AD** | Total Sites | 15866421 | 7132420 | 7589626 |
| | Total short variants (<10 bp) | 15555095 | 6057649 | 6031724 |
| | Variants in >= 5% of patients | 8517238 | 2002981 | 2000316 |
| | abs($\Delta AF_{alt}$) >= 0.1 | 2131794 | 387076 | 386650 |
| | $\Delta Wi_{INT}$ >= 0 | 179084 | 33469 | 33429 |
| | ref_prob > 0.5 & alt_prob <= 0.5 | 9483 | 5392 | 5389 |
| | ref_prob > 0.7 & alt_prob <= 0.3 And Exclude M/mAF<0.05 | 3692 | 2161 | 2159 |
| | LOD >= 13 (Valley inflection) | 2783 | 1721 | 1720 |
| **SCC** | Total Sites | 14294688 | 5214003 | 5686700 |
| | Total short variants (<10 bp) | 13850252 | 4383867 | 4369786 |
| | Variants in >= 5% of patients | 8571198 | 2020941 | 2018204 |
| | abs($\Delta AF_{alt}$) >= 0.1 | 1209636 | 220587 | 220154 |
| | $\Delta Wi_{INT}$ >= 0 | 139193 | 31723 | 31530 |
| | ref_prob > 0.5 & alt_prob <= 0.5 | 8185 | 4943 | 4930 |
| | ref_prob > 0.7 & alt_prob <= 0.3 And Exclude M/mAF<0.05 | 3709 | 2295 | 2288 |
| | LOD >= 16 (Valley inflection) | 2304 | 1578 | 1572 |

**B**

| Cancer: | Thresholds | Significant Number of across TFBS + Splice Region | | |
|---|---|---|---|---|
| | | **Unique Functional Site** | **TCGA Variants** | **dbSNP IDs** |
| | Total Sites | 16599737 | 9500957 | 9876505 |
| | Total short variants (<10 bp) | 16431811 | 8080612 | 8034254 |
| | Variants in >= 5% of patients | 8459283 | 2001368 | 1998783 |
| | abs($\Delta AF_{alt}$) >= 0.1 | 827197 | 152884 | 152443 |
| | $\Delta Wi_{INT}$ >= 0 | 81830 | 22796 | 22541 |
| **AD** | ref_prob > 0.5 & alt_prob <= 0.5 | 5156 | 3354 | 3335 |
| | ref_prob > 0.7 & alt_prob <= 0.3<br>And Exclude M/mAF<0.05<br>*<u>(Common variants kept)</u>* | 2327 | 1506 | 1490 |
| | LOD >= 17 (Valley inflection) | 1363 | 965 | 955 |
| | Total Sites | 16200808 | 8628213 | 8833051 |
| | Total short variants (<10 bp) | 15971353 | 6949822 | 6914833 |
| | Variants in >= 5% patients &<br>ref and alt match b/w dbSNP, cohort | 8515880 | 2014785 | 2012017 |
| | abs($\Delta AF_{alt}$) >= 0.1 | 1084527 | 199095 | 198651 |
| **SCC** | $\Delta Wi_{INT}$ >= 0 | 90377 | 24772 | 24550 |
| | ref_prob > 0.5 & alt_prob <= 0.5 | 5650 | 3689 | 3673 |
| | ref_prob > 0.7 & alt_prob <= 0.3<br>And Exclude M/mAF<0.05 | 2353 | 1524 | 1511 |
| | LOD >= 17 (Valley inflection) | 1391 | 995 | 985 |

**Table S4:** The table summarizes overlapping variants identified between cohorts, highlighting shared mutational features that may contribute to common biological mechanisms/ risk across patient groups.

| | Cancer: | Cohort-wise significant variants | Significant Number of across TFBS + Splice Region | | |
|---|---|---|---|---|---|
| | | | Unique Functional Site | # Variants | dbSNP IDs |
| All samples analyzed together | AD | TCGA | 1363 | 965 | 955 |
| | | CPTAC | 2783 | 1721 | 1720 |
| | | TCGA ∩ CPTAC | 334 | 246 | 245 |
| | SCC | TCGA | 1391 | 995 | 985 |
| | | CPTAC | 2304 | 1578 | 1572 |
| | | TCGA ∩ CPTAC | 826 | 604 | 600 |
| Smoking history-based separation | AD Smoker | TCGA | 2309 | 1598 | 1593 |
| | | CPTAC | 8352 | 5483 | 5481 |
| | | TCGA ∩ CPTAC | 820 | 590 | 588 |
| | SCC Smoker | TCGA | 2365 | 1625 | 1620 |
| | | CPTAC | 8766 | 5703 | 5704 |
| | | TCGA ∩ CPTAC | 982 | 680 | 681 |
| | AD NonSmoker | TCGA | 4830 | 3092 | 3096 |
| | | CPTAC | 18266 | 11789 | 11788 |
| | | TCGA ∩ CPTAC | 765 | 512 | 512 |

**Table S5:** Summary of genetic variants used as input features for the PGViS score. Variants were selected and incorporated into the patient-level bag of mutations according to analysis context: smoker-specific, non-smoker-specific, and pooled cohorts. *"All" denotes analysis without dividing samples by clinical variables.

| Dataset | Cancer (#Patients) | Significant Number of across TFBS + Splice Region | | |
|---|---|---|---|---|
| | | Unique Functional Site | Variants | dbSNP IDs |
| TCGA | AD Smoker (362) | 2309 | 1598 | 1593 |
| | AD Non-Smoker (61) | 4830 | 3092 | 3096 |
| | SCC Smoker (301) | 2365 | 1625 | 1620 |
| CPTAC | AD Smoker (133) | 8352 | 5483 | 5481 |
| | AD Non-Smoker (87) | 18266 | 11789 | 11788 |
| | SCC Smoker (84) | 8766 | 5703 | 5704 |
| CPTAC | All AD (227) | 2783 | 1721 | 1720 |
| | All SCC (109) | 2304 | 1578 | 1572 |

| | | | | |
|---|---|---|---|---|
| TCGA | All AD (434) | 1363 | 965 | 955 |
| | All SCC (329) | 1391 | 995 | 985 |

Note: *All is analysis without dividing samples based on clinical variables.

**Table S6:** Mapping for significant mutations common to both TCGA & CPTAC cohorts for the European ancestry. These were used to find smoking-dependent & -independent scores which was combined to generate PGViS for every patient. rGREAT: used to map the functional sites to nearest genes.

| Dataset (EUR) | Cancer | Type (# patients) | Germline mutations | dbSNP ID | Functional sites | Unique TF name | Unique Genes |
|---|---|---|---|---|---|---|---|
| CPTAC | AD | NonSmoker (12) | 480 | 480 | 708 | 228 | 320 |
| | | Smoker (74) | 487 | 487 | 658 | 220 | 225 |
| | | All* (89) | 181 | 180 | 233 | 132 | 98 |
| | SCC | Smoker (63) | 567 | 567 | 792 | 231 | 313 |
| | | All* (70) | 449 | 449 | 590 | 219 | 224 |
| TCGA | AD | NonSmoker (46) | 480 | 480 | 708 | 228 | 320 |
| | | Smoker (272) | 487 | 487 | 658 | 220 | 225 |
| | | All* (328) | 181 | 180 | 233 | 131 | 98 |
| | SCC | Smoker (222) | 567 | 567 | 792 | 231 | 313 |
| | | All* (240) | 449 | 449 | 590 | 218 | 224 |

*Note:*

1. Some of the genes may not map to any sites within the mutations so the number might drop from the initial tables.
2. *All is analysis without dividing samples based on clinical variables.
